# “It’s my job”: A qualitative study of the mediatization of science within the scientist-journalist relationship

**DOI:** 10.1101/2022.08.10.503486

**Authors:** Laura L. Moorhead, Alice Fleerackers, Lauren A. Maggio

**Author notes:** **Disclaimer**, The views expressed in this article are those of the authors and do not necessarily reflect the official policy or position of the Uniformed Services University of the Health Sciences, the Department of Defense, or the US Government.

## Abstract

Through 19 interviews with scientists, this study examines scientists’ use of media logic and their relationships with journalists using research as the focal point. The authors identified that the scientists shared a basic understanding of media logic classified in three patterns. Two patterns were previously identified by Olesk: 1) *adaption* (ability to explain research in a simple, engaging fashion but with a reactive approach to journalist interaction) and 2) *adoption* (proactively create and manage media interactions for strategic aims through a more active use of media logic). The other emerged as a new, third pattern, *affiliation* (enthusiastic contributors to journalists’ production practices and desire to engage in public outreach).

## Context

Research—typically in the form of peer-reviewed journal articles and preprints—acts as a basis of mutual interest for the scientist-journalist relationship. Journalists have long relied on these articles as a primary source for information in their reporting [Williams & Clifford, 2008; Veneu et al., 2008; Wihbey, 2017]. They connect with research articles in a variety of ways, notably through online databases, journals, and preprint servers, and media relations offices of universities, research organizations, and pharmaceutical companies [Amend & Secko, 2012; Fleerackers et al., 2021]. Journalists also reach out directly to scientists, asking for research articles and interviews [Fleerackers et al., 2021; Dijkstra et al., 2015]. Such approaches come with risks: a “loss of information diversity” through the repetition of information sources and the citation of the same scientists and research) and a science agenda overly influenced by academic institutions and scholarly and commercial publishers issuing press releases [Granado, 2011, p. 795]. These risks can be exacerbated by a mismatch in practices, norms, and values between scientists and journalists [Nguyen & Tran, 2019; Besley & Nisbet, 2013; Dunwoody & Ryan, 1985]. Scientists, for instance, may answer a journalist’s interview request or proactively share research with the goals of promoting their field, their research, or their institution, while journalists can be less concerned with these goals and more constrained by deadlines [Dijkstra et al., 2015; Peters, 1995].

Still, both journalists and scientists see science-media interactions as beneficial and the use of research as a shared touchstone [Dijkstra et al., 2015; Besley & Nisbet, 2013]. These symbiotic relationships between scientists and journalists [Lubens, 2015] may encourage an adaptation or adoption of practices between professions [Olesk, 2021]. Journalists’ roles have evolved [Fahy & Nisbet, 2011], with a move toward greater analysis and interpretation of research findings [Rensberger, 2009; Albæk, 2011]. Scientists, meanwhile, have moved toward, if not embraced, journalistic practices, goals, norms, and values in what they call the “mediatization of science”— “an increase in the orientation of science to its social context” [Peters et al., 2008, p. 72]. Mediatization can be understood as:

> the mutual relation between science and the mass media. It is based on the assumption that—due to the importance of the mass media in framing public opinion—there is an increasingly tighter coupling of science and the mass media [Franzen et al., 2012, p. 4-5].

This coupling has wide-ranging implications for science and society, shaping how scientific research is conducted [Weingart, 2012] and presented to the public [Peters, 2012]. As such, scholars have raised concerns that mediatization encourages a weakening of science’s autonomy [Weingart, 2012] through a bend toward “media logic,” which Altheide [2013] described as the form and formats of communication. Alongside these potential dangers of mediatization, however, scientists’ adoption of media logic could also facilitate interactions between journalists and scientists by providing the shared norms, practices, and expectations needed to effectively communicate [Carson, 2015].

To understand the implications of the growing mediatization of science [Bauer, 2012], scientists’ use of media logic must be considered alongside ongoing changes to the relationships between scientists and journalists. Recent studies suggest that scientists and journalists align in their motivations, particularly in their sense of shared public responsibility and push for responsible research [Olesk, 2021; Dijkstra et al., 2015]. Increasingly, journalists rely on interviews with scientists to legitimize their news frames and to facilitate a “dynamic interplay between journalist and researcher that will largely determine whether or not the journalist comes to see the event as sufficiently ‘significant’ and ‘interesting’ to warrant news coverage” [Albæk, 2011, p. 344]. The growing value placed on public visibility within the culture of science may also be increasing scientists’ reliance on journalists [Dunwoody, 1999; West & Bergstrom, 2021]. Dunwoody [1999] argued that we should expect to see journalists and scientists develop a “shared culture,” in which both groups *equally* contribute to the public portrayal of scientific evidence. In such a culture, scientists would no longer simply be passive sources of information but active partners in newswork—working alongside journalists to select, interpret, and communicate research evidence to society. While this affiliation between scientists and journalists may lead to smoother interactions and an easier reporting process, it may also challenge the watchdog role of journalists [Cormick, 2019; Schulson, 2016].

While mediatization has been extensively considered (and debated) [Weingart, 2012; Wihbey, 2017], it is unstudied in light of Covid-19 as an ongoing global health crisis with associated scientific controversies (i.e., efficacy of vaccines, mask mandates, use of preprints). Bucchi [1996] noted that scientists, when faced with controversy in their fields, work to address the public directly. Perhaps unsurprisingly, Covid-19 has encouraged scientists to do this through online and social media [Bhopal & Munro, 2021; Colavizza, 2021; Joubert, 2020], including through publishing models such as *The Conversation*, which partners scholars with journalists [authors, under review]. These evolutions in the ways scientists and journalists communicate reflect the kind of *post-normal science communication* (PNSC) [Brüggemann et al., 2020] that can take place when “facts [are] uncertain, values in dispute, stakes high and decisions urgent” [Funtowicz & Ravetz, 2020, p. 1]. In such contexts, journalists and scientists may come to share norms, practices, and goals, as the boundaries between the two fields blur and are renegotiated [Brüggemann et al., 2020]. This renegotiation—continuing throughout the pandemic and possibly expanding in an era of declining public trust in scientists and journalists [Kennedy et al., 2022]—will likely lead to the adoption of new norms and practices, which may, in turn, affect the nature of relationships between scientists and journalists.

## Objectives

This study, conducted during the pandemic and in the context of scientific debate, controversies, and political polarization [Dunwoody, 2020], aims to examine scientists’ use of media logic and the nature of their relationships with journalists. It does so using research (i.e., preprints and peer-reviewed journal articles) as the focal point for qualitative interviews, offering a view into this intersection of seemingly disparate professions as they negotiate the volatile waters of our global pandemic.

We apply the mediatization of science as our conceptual approach and adapt a framework by Olesk [2021] to evaluate the mediatization patterns of scientists in relation to journalists, making this one of few studies that have investigated science-journalist interactions using an explicit theoretical framework [Dijkstra et al., 2015]. We add to Olesk’s [2021] list of indicators, using scientist interactions with journalists to develop scientist personas—which Daston and Sibum [Daston & Sibum, 2003] called *cultural identities*—that might allow for a more nuanced understanding of scientists’ professional roles alongside their personal needs, experiences, behaviors, and goals in the scientist-journalist relationship. More specifically, we ask:

RQ1: What indicators can be used to expand and describe the mediatization patterns of scientists who engage with journalists?
RQ2: What scientist personas can be identified using these indicators?

## Methodology

This study is part of a larger research program that explores scientific research featured in the news. We focus primarily on scientists’ perspectives; however, our analysis was informed by journalists’ interviews (see Authors, 2021). We conducted the current study using qualitative description methodology [Sandelowski, 2010] guided by a constructivist paradigm [Sandelowski, 2010]. Constructivism assumes that participants devise the realities in which they engage. Through this lens we were able to better understand scientists’ motivations, views, and professional practices in relation to journalists.

### Recruitment

We recruited 19 scientists whose research had been mentioned in a news article. The lead author and a research assistant manually identified names of scientists who were quoted directly or whose research was mentioned or hyperlinked in a sample of 400 news articles from *The Guardian*, HealthDay, IFL Science, MedPage Today, News Medical, *New York Times, Popular Science*, and *Wired*.^1^ Each article mentioned at least one preprint or peer-reviewed research article; news articles were gathered during March and April 2021 (see Fleerackers et al., 2021, for detailed data collection process). This study was exempted from further review by two university ethics boards [institution names and REB numbers anonymized for peer review].

### Interviews

We designed our semi-structured interview protocol^2^ using the literature and our experience as journalists and research scientists. The first portion of the protocol included general questions about scientists’ use of research and experience working with journalists; the second portion was a talk-aloud in which they described their actual experience in the reporting of a science news article drawn from our sample. This approach allowed scientists to say what they typically did (first portion of interview) and then explain what they actually did for a particular story (second portion). Recruitment and interviews occurred between September-January 2022. After 15 interviews, we began to discuss the potential of reaching an adequate level of information power base that would enable us to meet our research aims [Malterud et al., 2016]. After 19 interviews, we agreed that we had reached an adequate level. Interviews, which lasted up to 60 minutes, were conducted and recorded via Zoom and were transcribed and de-identified for analysis.

### Analysis

We used framework analysis [Ritchie et al., 2013], which accommodates multidisciplinary research teams and thematic analysis of semi-structured interview transcripts [Gale et al., 2013]. The framework allowed us to compare and contrast data across cases, as well as within individual cases (i.e., individual scientists), and to identify first patterns and then personas of mediatized scientists. We independently read and coded each transcript, using a mix of deductive coding based on Olesk’s [2021] existing typology of mediatized scientists) and inductive coding (based on emergent patterns in the data). We coded instances of scientists presenting indicators of media logic in five dimensions (see Table 1). Throughout the coding, the three authors (a professor of medicine and two former journalists now working as academic researchers) met multiple times virtually to reflect on the analysis. In these conversations, we recognized and discussed how our backgrounds and experiences facilitated our ability to be reflexive in our examination of the transcripts from the perspective of both professions.

**Table 1.** Framework used to analyze mediatization of scientists. Codes for Adaption and Adoption were drawn from Olesk’s [2021] original typology, codes for Affiliation were developed inductively.

| Dimension | Definition | Adaption of media logic [Olesk, 2021] | Adoption of media logic [Olesk, 2021] | Affiliation of media logic (inductively derived) |
| --- | --- | --- | --- | --- |
| Communication as a responsibility | See public communication as part of professional responsibility. | See it as important but secondary to their scientific work. | See it equally important to their scientific work. | See it as a shared responsibility with journalists. |
| Awareness of media logic | Express awareness of media logic and feel confident using journalistic news style. | Able to explain their work in simple terms and feel confident giving interviews. Criticize journalists' routines. | Able to understand and accept journalists' work logic, and express themselves in journalistic news style. | Able to appreciate and contribute to content production practices used by journalists. |
| Mastering media logic | Confident in mastering media logic and using it to trigger media coverage or introduce relevant angles. | Not familiar with news production practices; write occasional press releases; otherwise do not initiate media coverage. | Contact journalists proactively and “sell” stories and angles to them. | Articulate and share content production practices used by journalists. |
| Purposeful use of media | See media as a tool for achieving scientific or non-scientific aims. | See media coverage as benefitting the current project (getting attention, increasing awareness about an issue, etc.). | Have more strategic aims (wider benefits to science, economy, etc.) and think in terms of *target groups* and *messages*. | See participation in media coverage as a partnership with journalists with a desire for high impact (contribute to public health, society) and think in terms of *behavior change* for public. |
| Institutionalization of communication activities | Communication activities in the research group/organization have been institutionalized within the professional activities of the scientist. | Perform communication activities on *ad hoc* basis. | Conduct communication activities systematically and follow a strategic plan, integrating public communication into the professional activities of the scientist. | Has internalized communication activities, integrating them into both professional and personal activities (e.g., tweet, personal blog). |

We used a spreadsheet to generate a matrix, including references to illustrative quotations. Charting involved summarizing the data by category to create a holistic impression of what each scientist said [Miles et al., 2018]. For each scientist we created a *user profile* with demographic information and categorized each scientist’s orientation to media logic using the adapted Olesk [2021] framework (see Table 1). Then, we created *personas* (i.e., “super-typical” representations of scientists) by grouping user profiles based on demographic and mediatization patterns [LeRouge et al., 2013].

## Results

All 19 participant scientists shared at least a basic understanding of media logic; yet our analysis revealed three patterns in their mediatization. Two of these patterns were previously identified by Olesk [2021]: 1) *adaption of media logic* (ability to explain research in a simple, engaging fashion but with a reactive, rather than proactive, approach to journalist interactions) and 2) *adoption of media logic* (proactively create and manage media interactions for strategic aims and through the more active use of media logic). We identified a new, third pattern, *affiliation of media logic*, through early reading and coding of the transcripts.

### Patterns of mediatization

#### Adaption of media logic

A minority of scientists expressed an awareness and basic mastery of journalistic norms, values, and practices but took a reactive approach to communication activities characteristic of adaption of media logic. These scientists’ interactions with journalists were typically mediated by their institution’s communications group or a journal’s press department. While these scientists could articulate the process of working with the media, they did not necessarily experience it firsthand, often relying on others to write their quotations and public-facing research descriptions. They did not prioritize media outreach or see their relationships with journalists as something they needed to maintain or improve.

Adaptive scientists generally viewed the role of communication professionals as helpful—a shield from the risks of working with the news media. If a journalist reached out to them directly, they typically reported seeking help in responding from their institution’s communications group. As one scientist explained,

> We have people here who write drafts, and we’ll go back and forth and make sure that the science and, you know, the communication is as accurate as it can be. And then, they do the press release, and then news outlets will take that up. [Sci_13]

As adaptive scientists’ understanding of the journalistic process was typically framed from the perspective of an institution or journal, they could be flummoxed when a journalist deviated from this idealized process. For example, adaptive scientists were often frustrated if journalists did not circle back with their quotations and interview content for approval or did not exclusively contact first or second authors.

Adaptive scientists also viewed journalism as a way to promote one’s work, rather than to promote research as a societal good. Several resisted reaching out to journalists, with one saying:

> I do not…contact journalists to send them my work. I don’t know, it feels like—for some reason—it feels tacky to do that, but maybe I should do it more often? [Sci_11]

Yet, that scientist also acknowledged that peers had different, albeit still promotional, approaches:

> We have this paper that’s currently in review, and the first author, who is currently looking for a new job, was very excited and started posting the preprint, started showing it around. Journalists started contacting him to interview…and, actually, a piece came out at some point. I was not angry, because I can understand why he did that. [Sci_11]

Adaptive scientists also seemed to share a lack of confidence in journalists’ ability to understand research:

> Unfortunately, it just seems really, really, really unlikely to me that a journalist can look at a preprint or an article, something in arXiv, and make any sense out of it, and make any judgment about correctness or importance, or anything like that. [Sci_08]

#### Adoption of media logic

Adoption of media logic was a more common pattern among the scientists, characterized by ambivalence about the controlled forms of media outreach laid out by communication groups at their institutions and target journals. On the one hand, adoptive scientists said they appreciated the efficiency of this approach; they put their trust in communications professionals, thankful that someone else could handle the influx of media requests and help them navigate the media system. On the other hand, adoptive scientists were sometimes frustrated that this approach gave communications professionals ultimate control over their public communication. They lamented how communications professionals organized and “triaged” media interviews, deciding which requests to prioritize and which to pass over; determined which papers to promote actively; enforced limits on what scientists could and could not discuss on the record; and prepared press releases with ready-made author “quotes” for scientists to review and approve. Unlike their adaptive peers, adoptive scientists also revealed a sophisticated understanding of the outcomes of media coverage, which they leveraged to advance their institution’s brand and reputation, recruit faculty and students, and procure funding. Adoptive scientists better recognized that journalists operated independently—outside the controlled, if not idealized, realm of an institution’s communications group or a journal’s press department.

Adoptive scientists also proactively created and managed media interactions, stating that they “always respond” or “try to respond to all” journalist inquiries [Sci_12, Sci_19]. These scientists considered working with journalists as part of their professional role, even if the effort fell outside of their formal work description. One scientist put it simply: “It’s my job” [Sci_19]. Another scientist explained his need to “always respond” in the context of journalists’ reliance on experts for accuracy:

> The last thing I want is for a journalist to write a paper about our work or about anybody’s work without talking to experts, so I’m totally available… we want it to be presented in the best, correct scientific light.” [Sci_12]

Oftentimes, adoptive scientists leveraged multiple ways to encourage media coverage. One scientist said, “I know journalists cover scientific conferences,” adding that he responded to a journalist at a recent conference and “ended up exchanging emails” as the journalist “prepared the piece” [Sci_18]. That same scientist also reached out directly to journalists through a range of media, intertwining personal and professional realms:

> I think leveraging all of those resources—social media through your own personal or institutional account, but also using media outlets virtually or in print—could be very beneficial for scientists. [Sci_18]

As demonstrated above, adoptive scientists often employed language suggesting the “use” of journalists and the media to achieve their goals. However, they also expressed frustration at the professions’ differing practices. One scientist explained,

> Sometimes it can be challenging talking to journalists, and there’s just different norms about attribution and citation and stuff like that in journalism versus academia. [Sci_14]

Scientists’ lived experiences interacting with journalists did not always align with their more abstract, big picture reflections on their relationships to media logic. Even scientists who held journalists in high esteem overall could recall negative interactions (e.g., being misquoted, being asked unexpected or inappropriate interview questions). These negative experiences often elicited critical perceptions of journalists that were most in line with an adaptive orientation. This was the case for Sci_05, for example, who recounted a live radio interview in which she was not addressed by name but instead referred to as the “pretty mumps lady.”

In most cases, however, specific experiences working with journalists did not appear to fundamentally change adoptive scientists’ underlying orientation to media logic; instead, they elicited a more measured approach to media interactions, particularly when accepting interviews from unknown journalists or those from outlets perceived to be less trustworthy. For instance, several scientists preferred national over local media and legacy print publications over radio and broadcast outlets, believing that those publications produced higher quality journalism.

Despite negative experiences with *specific* journalists, these scientists expressed deep, if not grudging, respect and gratitude when speaking about journalists in general. As one senior scientist commented:

> It’s rare for me to talk to a journalist who doesn’t show a very intelligent understanding of the field. They’ve done their homework…. If they’re confused about something after the interview, they call me back and get clarification about something. [Sci_12]

#### Affiliation of media logic

Adding to Olesk’s [2021] framework, we identified a third pattern of mediatization—affiliation of media logic—that differed from the other patterns in important ways. Affiliative scientists demonstrated greater contribution to the content production practices of journalists than either adaptive or adoptive scientists. They cared deeply about public outreach, unlike adaptive scientists, who saw media communication as secondary to their research. In this sense, they resembled adoptive scientists, who saw communication as equally important to their other professional duties. Yet, affiliates expressed greater appreciation for journalists’ unique abilities and used their awareness and mastery of media logic to support, rather than control, journalists’ work. Finally, although these scientists pursued communication with journalists in a goal-oriented fashion that resembled adoption of media logic, they did so with broader, societal goals in mind (e.g., reduce Covid-19 transmission, promote vaccine safety, etc.).

At the core of the affiliation of media logic pattern was a sense of collaboration. These scientists partnered with or helped journalists by articulating research in simplified narratives, providing critique and context about other scientists’ studies, and inspiring news frames and story ideas. They also employed characteristics of content production used by journalists, such as an awareness of a story’s timeliness and the need for it to be both interesting and relevant to the public. These scientists displayed a more purposeful use of media. For instance, one scientist said, “For me, I see the press as an ally in terms of helping disseminate information and being very committed to doing that accurately and fairly” (Sci_19). Many enjoyed talking with journalists and some believed their research benefited from the conversations. Several had fostered long-term relationships with journalists, who would occasionally call on them seeking comments about new studies in their area of expertise. These scientists also recognized journalists’ unique skills in explaining difficult or technical research to the layperson. As one participant explained,

> There has been a lot of news or information on the virus that was not precise […] and that’s really a problem. From that point of view, the scientists—the ones that are really working on things—they should really help the journalists provide reliable information. [Sci_01]

For affiliative scientists, interactions with journalists were motivated by a personal mandate to communicate science, beyond any expectation to follow media relations protocols laid out by research institutions or journals. These scientists integrated public outreach into both their professional and personal activities, for example, by sharing their research on social media, personal websites or blogs, or through articles contributed to “research amplifier” platforms [Osman & Cunningham, 2020] such as *The Conversation*.

### Factors intersecting with mediatization

While the adaption, adoption, and affiliation patterns appear clear cut, most scientists did not consistently follow a single pattern for all five dimensions of mediatization. Instead, most expressed different patterns depending on the dimension, expressing, for example, an affiliative pattern for the *Communication as a responsibility* and *Purposeful use of media* dimensions, but an adaptive pattern for the *Awareness, Mastering*, and *Institutionalization of media logic* dimensions. We explored these variations further while developing scientist personas, reflecting on their relationships with other aspects of participants’ profiles. We found that three interconnected factors—career status, journal pressures, and institutional context—intersected with scientist mediatization patterns to shape their interactions with journalists.

#### Career stage

Among participants it was clear that early career scientists experienced risks that more established scientists did not. Keenly aware of the embargo policies at their target journals, and the importance of publishing in “high impact” venues, these untenured and early-career researchers (ECRs) often hesitated to fulfill journalists’ requests. This barrier was most obvious in the case of unpublished data and preprints, which they did not want to discuss with journalists for fear of jeopardizing future publication opportunities. For example, one ECR recounted a time that they had been approached by a journalist with a request for unpublished data. While the scientist wanted to contribute, and felt that their evidence would have enhanced the journalist’s story, they were unable to share the data because “that’s a huge career issue for me if I just kind of give it up” as “a lot of journals won’t let you submit if you shared your information or shared your data elsewhere” [Sci_16].

ECR status amplified not only professional risks but also potential benefits of interacting with journalists. When it came to peer-reviewed research, ECRs stressed that media coverage “does help our careers quite a bit with the tenure process” [Sci_09]. Some established scientists similarly acknowledged that this attention “can be seen to be sort of good for the CV/career” [Sci_02] as “the paper you published is somehow more important than if you don’t have press” [Sci_01]. Yet, more senior scientists described these career rewards as more of an added benefit than a major motivation for working with journalists, possibly because—as tenured researchers—they had already proven their value at their institution.

#### Journal pressures

The need to please journals was an important force shaping scientists’ interactions with journalists. This pressure meant that both fears and potential benefits associated with media attention were often amplified when submitting research to “high impact” journals, which scientists believed were not only more valued by their tenure committees but also by their institutional communication groups. As one researcher recounted of her time as a grad student:

> I was at, you know, a big R1 university, and the culture was sort of if your paper wasn’t in *Science* or *Nature*—or maybe *PNAS*—like, you did not tell the press department. Like, they only cared about high-impact articles. [Sci_10]

Beyond implicit pressures related to the journal publishing system, journals directly shaped scientists’ interactions with journalists by setting embargoes, preparing and publicizing press releases, and promoting new studies. Again, “high impact” journals appeared to play an outsized role. As one participant explained:

> …they are very keen on broader dissemination. So if you publish in the high-impact journals, you know, I think the aim is that it gets out to a wider audience, by default. And they have a very active kind of media division. [Sci_02]

#### Institutional context

Finally, scientists’ institutional context informed whether and how they engaged with journalists. Institutions directly influenced interactions with journalists by preparing press releases, pitching media coverage, facilitating interviews, and more. Some also had strict policies controlling how or whether employees could engage with journalists. As one scientist explained, these policies sometimes acted as a barrier to communication:

> …because it’s a US government organization, we have to be really careful about not appearing to endorse products or things like that. So sometimes explaining what we’ve done is difficult. [Sci_06]

Institutions also implicitly affected interactions by communicating norms and expectations about what kind of media attention, if any, was considered acceptable and valuable. Some actively encouraged and rewarded media outreach (e.g., those with active communications groups, media training opportunities, public outreach mandates), while others were less enthusiastic. For one scientist, a lack of alignment between their personal communication goals and those of their institutional context encouraged a job change. This scientist explained:

> If you can get your work into open-access journals or you work for an institution that’s willing to pay the open-access fees, then I think academic journals can be a reasonable vehicle of influence, as long as the turnaround is quick. But my experience is that it takes a while […] so I took a job with [another institution], which produces research to inform law and public policy, in part because I felt like I wanted my time to be meaningfully spent […] We’ve got a great communications director, and our work is often cited by the press, more so than when I worked at universities. [Sci_19]

### Scientist personas

By combining the scientists’ accounts through Olesk’s framework and three interconnected factors, we developed four personas that allowed for an intertwining of dimensions and, ultimately, a more complex and nuanced understanding of scientists’ professional roles alongside their personal needs, experiences, behaviors, and goals in the scientist-journalist relationship. These fictional personas represent different scientist types that interact with journalists. Displayed by their profiles below, the personas were the (1) Constrained Communicator, (2) Ambivalent Media Source, (3) Strategist, and (4) Media Enthusiast.

#### Constrained Communicator

**Figure 1.**
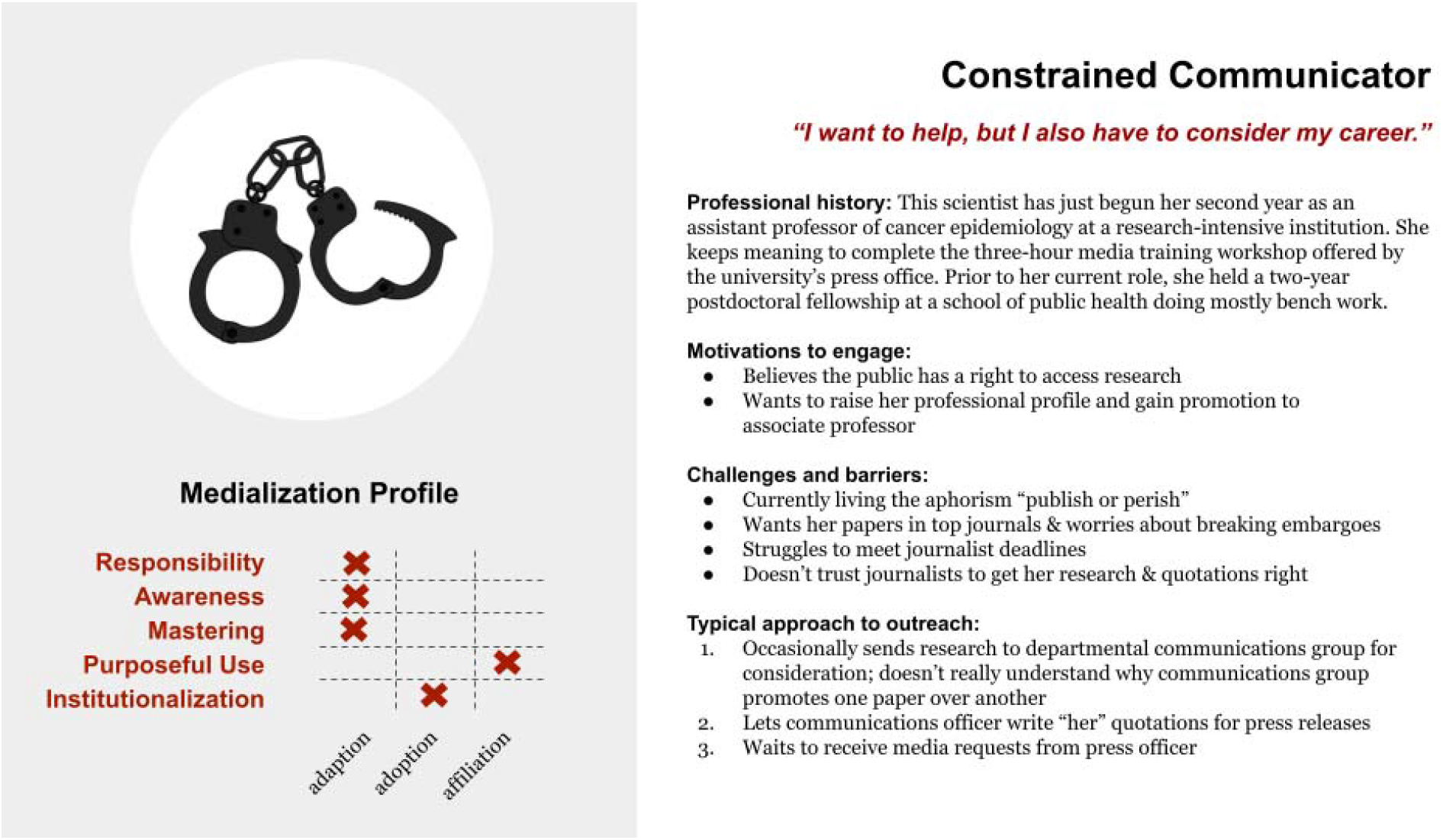
Persona representing the Constrained Communicator.

While most scientists reported some level of pressure or control from their institution or target journals, the Constrained Communicator expressed the greatest frustration. Typically, this scientist was either an ECR focused on academic promotion or a more senior researcher working for a large nonprofit or government organization. The ECR found the general publishing and research promotion process—from journal embargos to institutional and journal press releases— frustrating and out of their control. As one scientist explained,

> [The press release] was from the journal. They wrote one. And we actually didn’t have a lot of say in how that was written. I remember because we were not super happy with it, but they said we should just only comment…. They made it very clear that we were not supposed to change anything. It was odd. [Sci_03]

Senior scientists were more accepting of institutional processes, but felt those elements could work against a mandate of science: to share timely, understandable scientific knowledge with the public.

Both types of Constrained Communicator viewed their plight as part of a larger promotion system outside their control. They followed the lead of press officers, hewed closely to dictums of top journals, and resisted sharing data and papers prior to peer-reviewed publication. Contact with journalists was highly mediated by their institution; their views of journalists were framed by communication professionals.

#### Ambivalent Media Source

**Figure 2.**
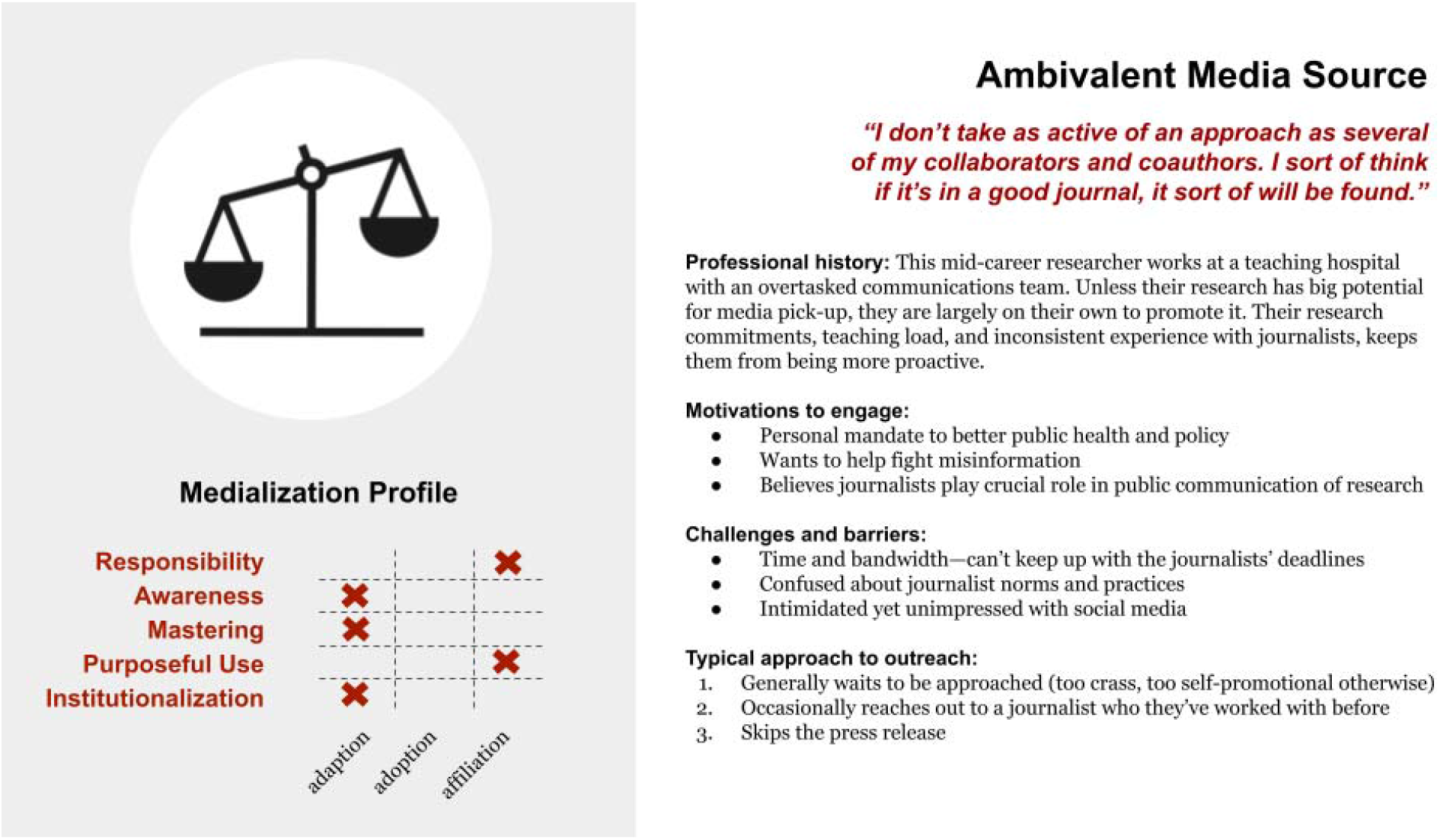
Persona representing the Ambivalent Media Source.

Typically a mid-career scientist, the Ambivalent Media Source expressed mixed feelings regarding their interactions with journalists. While they believed journalists could be crucial in “translating” esoteric research, they also worried about journalists’ accuracy. This scientist bemoaned losing control of their research:

> Once it is published, everybody can read the article, and it is not our thing. I mean it’s something public. And it’s okay, but sometimes when things are not accurate it’s a bit sad. [Sci_07]

They were also more pessimistic about the scientist-journalist relationship, occasionally speaking of the two professions as misaligned in goals, norms, and professional practices:

> Maybe I’m too cynical, but it feels like…. They’re covering a particular issue for a reason; sometimes it could be that they’re genuinely interested in learning about new developments in a particular field. But I think many times, journalists—they already know the content… of the piece they’re going to write. [Sci_18]

Perhaps as a result, the Ambivalent Media Source rarely approached journalists, recognizing that they would need to commit time and energy—which they did not have—to communicate. Additionally, there were no guarantees that their efforts would pay off:

> Sometimes, I’ll spend an hour or so talking to a journalist, and then they’ll use a lot of the stuff I told them, and not mention that I was the one who told them or not link to any of the papers. [Sci_14]

While the Ambivalent Media Source occasionally had direct contact with journalists, it was reactive (i.e., “I always call them back”) and their mediatization was piecemeal, with a limited understanding of journalistic practice. One scientist said:

> We get contacted by journalists and then even if you ask them, ‘Can you send me a link when the piece is out,’ they rarely do it. I don’t know if it’s some rule to not do it, or if they just forget or don’t care. [Sci_11]

#### Strategist

**Figure 3.**
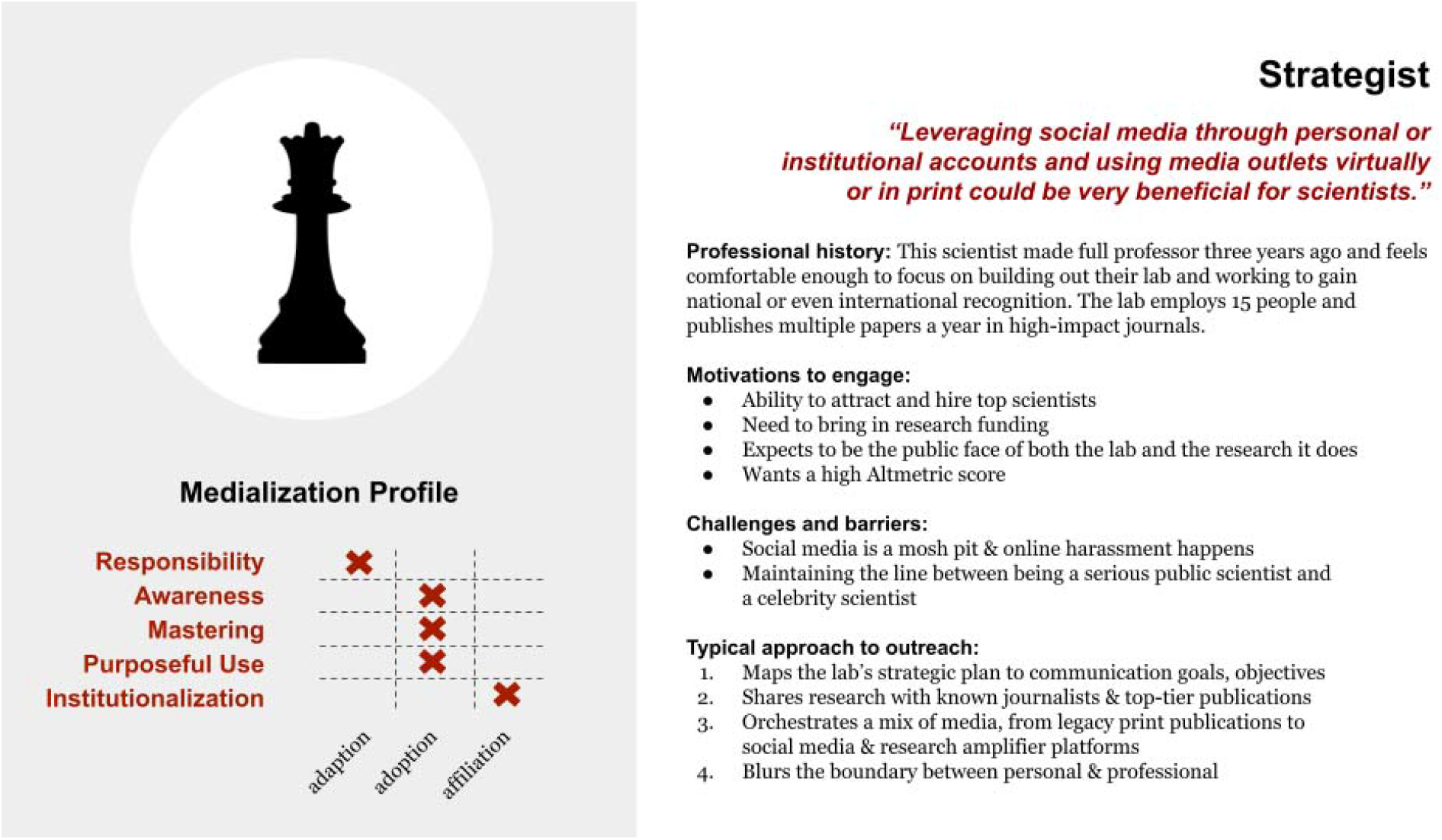
Persona representing the Strategist.

This seasoned scientist was a strategic marketer and recognized their efforts as crucial for gaining talent and funding. Their lab and research were well established and did not require constant oversight, allowing time and space to develop plans for promoting their work. The Strategist saw media coverage as a powerful tool for advancing their career, findings, and field. They worried less about being seen as crass or “tacky” and were comfortable using mediatized, commercial language:

> Getting some recognition… is motivating… exposing the public to some of the nice things that you are doing—that’s one of the greatest recruiting tools for science and engineering. [Sci_09]

From their view, journalists and communication professionals can—and should—be managed. The Strategist was selective about the media they shared their research with, favoring journalists at major legacy publications such as the *Atlantic*, BBC, *Guardian*, *New York Times*, or *Washington Post*.

Highly mediatized, the Strategist used many journalistic tools and approaches to orchestrate media coverage and dedicated considerable time to planning interactions with journalists.

#### Media Enthusiast

**Figure 4.**
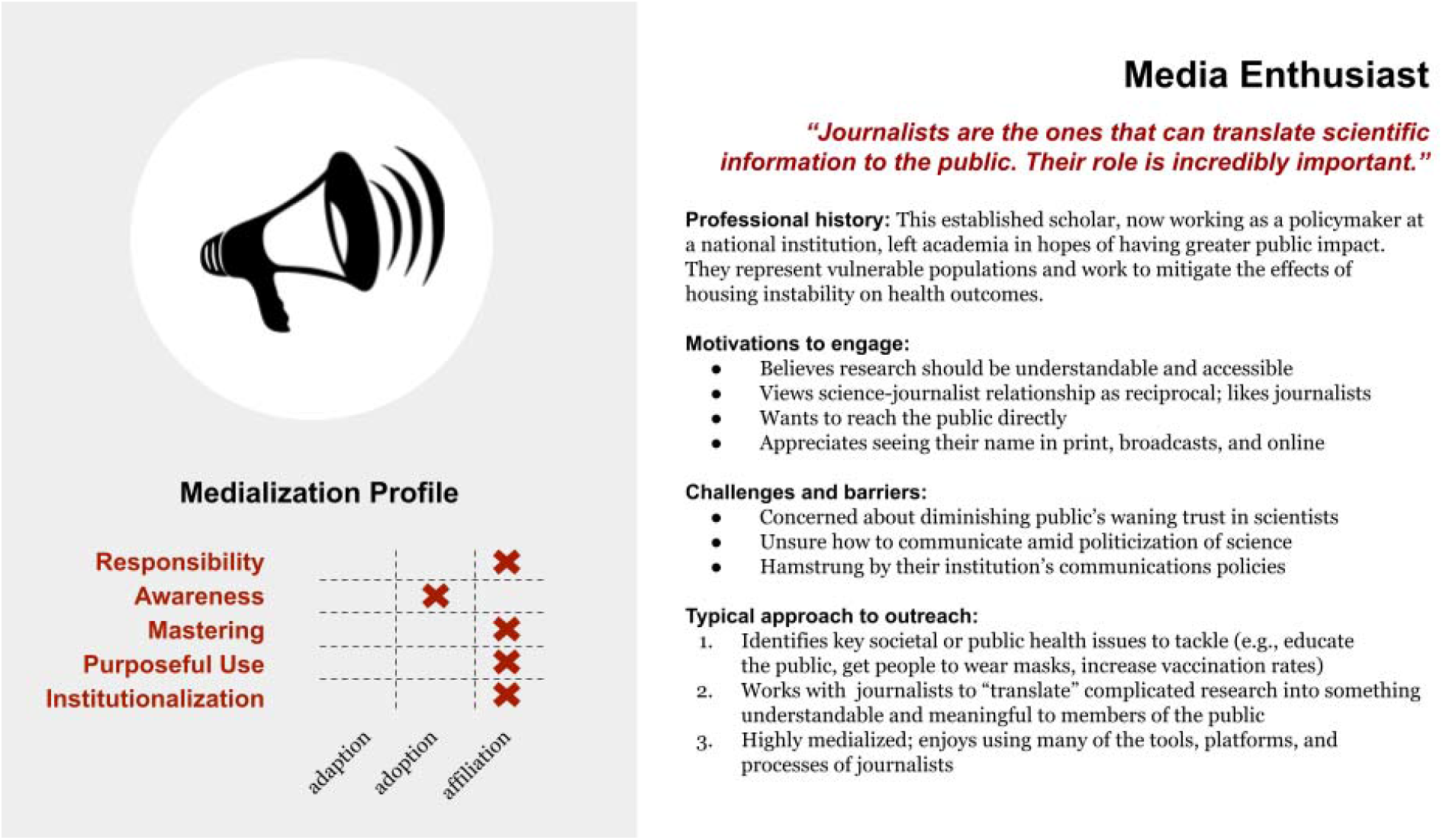
Persona representing the Media Enthusiast.

This scientist genuinely liked working with journalists and saw their efforts to do so as a collaboration. The Media Enthusiast viewed journalists as a key way to share knowledge and encourage change in society. They were mediatized enough to be strategic but were driven by a desire to share science, rather than promote themselves or their institutions. As one Media Enthusiast put it:

> [Getting research to the public]—it’s huge. I mean, most of my work is publicly funded, so I have a mandate or mission to share the work with the public… journalists are and news organizations are super, super important in that process…. They are the best way for us to get the word out. [Si_10]

The Media Enthusiast had often left—or was teetering on leaving—academia for an environment where they believed they would have greater public impact.

Highly mediatized and curious, the Media Enthusiast was likely to join a journalist for lunch at a conference or exchange email messages about a topic of interest. They brainstormed with journalists and shared story ideas. They regularly tweeted, looked for opportunities to build their online following, and enjoyed publishing work on research amplifier platforms such as *The Conversation*, where they could partner more formally with journalists.

## Discussion

This research examines the mediatization of science holistically, exploring how scientists’ professional context works alongside their internalization of media logic to shape interactions with journalists. Our findings offer a comprehensive and updated understanding of mediatization, demonstrating how factors such as career stage, pressures from journals, and institutional context can intersect with a scientists’ wider communication goals to influence whether and how they engage with journalists. We also highlight a partnership-type “affiliation” orientation of scientists to journalists that is characterized by collaboration, shared interests, goals, and efforts. In doing so, we make several empirical, practical, and methodological contributions.

Empirically, the affiliation pattern expressed by many scientists in this study diverges from previous research suggesting antagonistic relationships between scientists and journalists [MacNamara, 2014], but supports recent studies suggesting such relationships are generally positive and mutually beneficial [Peters et al., 2008; Dijkstra et al., 2015]. The dominance of this affiliative pattern also aligns with Dunwoody’s [1999] prediction—made two decades ago—that a “shared culture” would eventually emerge between scientists and journalists, in which the two sets of actors would equally contribute to the public communication of science. What this affiliation orientation means for science, journalism, and the public is unclear. On the one hand, mutually supportive relationships between scientists and journalists could support high quality, evidence-based science media coverage—particularly given that this orientation is characterized by goals of improving public wellbeing and maximizing societal benefits. On the other, the affiliation orientation could signal a further breakdown of the autonomy of science [Weingart, 2012] and of journalism [Schulson, 2016].

Our study sheds light on the interconnected dimensions and roles that personal, institutional, and systemic factors can play in the mediatization of science. In developing our personas, scientists’ career stage, institutional contexts, and pressures from journals emerged as important forces shaping the nature of their relationships with journalists. This echoes findings by Calice et al. [2022] that institutional factors, particularly in regard to tenure and promotion, are crucial in whether or not a scientist will engage with the public. Our study offers a view into how communication professionals at both academic institutions and scholarly journals implicitly and explicitly influence scientists’ participation in that competition, with implications for how scientists and journalists work together in the public communication of science. In particular, our study reveals the often overlooked role that scholarly publishing plays in whether and how scientists participate in public engagement. Findings suggest that journals may, in fact, have their own form of mediatization, in which scientists bend toward their norms and practices more than those of journalists. At times, the pressures to publish in high-impact journals discouraged even the most affiliative scientists from discussing their research with journalists before it had been peer reviewed and published. At others, journals facilitated media outreach by preparing press releases, introducing embargoes that allowed more time for scientist-journalist interaction, and arranging interviews to promote new publications. Such facilitation was typically welcomed by scientists, but allowed for a high level of control, from dictating scientist quotations to directing the news cycle of science and potentially narrowing information sources by favoring particular journalists and media organizations [Granado, 2011]. The role of the journal system as both an enabler and obstacle in the public communication of science warrants further research, particularly as embargoes and press releases influence the work of journalists and, ultimately, what knowledge is shared with the general public [Sumner et al., 2016; Taylor et al., 2015].

Practically, our findings could help address concerns that scientists need more than tactical skills (e.g., speaking and writing clearly, fostering dialogue, telling stories) for engaging the public with their research [Besley, 2020; Cooke et al., 2017]. Besley pointed out that “most communication experts within the scientific community work for organizations where the primary goals are about helping the organization, rather than advancing the overall scientific enterprise” [Besley, 2020, p. 158]. Our findings also point to this concern and extend it beyond what Besley called “the health and welfare of science” to the health and welfare of society. Personas from this study could be used to develop guidelines for supporting scientists of different institutions, career stages, and mediatization patterns to engage in strategic science communication for the benefit of society. For instance, our findings support calls to support faculty members in pursuing meaningful public engagement through changes to review, tenure, and promotion guidelines (e.g., Calice et al., 2022; Alperin et al., 2019]. Our findings could also help communications professionals at institutions and journals adapt their policies and systems to ensure they enable, rather than inhibit, accessible, impactful, and societally beneficial media coverage of research.

Our study makes several methodological contributions. It introduces a novel methodology that integrates framework analysis and persona development to provide theoretical and practical insights. It also highlights the value of using methods such as talk-alouds or reconstructive interviews to anchor discussions of relatively abstract topics to real-world practices (cf. Barnoy & Reich, 2019, 2022]. For example, the scientists sometimes described their relationships and practices differently when answering general, open-ended questions than when discussing specific news stories during the “talk-aloud” portion of their interviews. We encourage scholars to integrate the two elicitation approaches, as the tensions between the general and the specific that emerged during the interviews added a richness and complexity to the data that allowed us to answer our research questions with greater depth and nuance than would have been possible using either interview method alone.

This study must be considered in light of its limitations. We conducted research at a time of relative stability during the pandemic; the initial vaccine rollout had been completed and boosters were being administered in the US, Canada, and the UK, where most of the scientists were based. It is likely that the views in this paper would differ from those of scientists interviewed at the onset of the pandemic. Also, it is possible that this relatively high level of mediatization of participants is, in part, an artifact of when the interviews were conducted (i.e., during the COVID-19 pandemic). However, although a few scientists in our sample described increased or altered media relations as a result of the pandemic, the vast majority of scientists we interviewed did not describe altered media relations as a result of the pandemic. The timing allows for us to link the changing practices and norms of journalists to the changing (i.e. post-normal) communication context. All scientists had research mentioned in at least one article by a journalist working for a science publication. As such, these scientists may have had a higher degree of mediatization than scientists outside our study. Additionally, all publications in the data set were text-based (not multimedia), English only, and based in the Global North. Future research could expand outside these three categories.

## Conclusion

In conclusion, regardless of specific orientation to media logic, most, if not all, of our scientists can be described as relatively mediatized. All understood at least some of the norms, values, and practices of journalists and had been interviewed for news stories. Many also knew how to use their knowledge of media logic to pursue professional, institutional, or societal goals. This suggests an area ripe for future research in order to understand how to best support scientists and journalists in increasingly collaborating to sharing research with the public through the news media.

## Appendix A. Supplementary data (online)

**Table 1.** User profiles for interviewed scientists

| Scientist | Country | Experience | Institution (type) | Journal articles published in past year | Persona |
| --- | --- | --- | --- | --- | --- |
| Sci_01 | France | Senior | National research institution | 5 | Constrained Comm |
| Sci_02 | England | Senior | University | 12 | Media Enthusiast |
| Sci_03 | England | ECR | University | 4 | Constrained Comm |
| Sci_04 | U.S. | Senior | University | 50 | Media Enthusiast |
| Sci_05 | US | Senior | Local government | 4 | Strategist |
| Sci_06 | US | Senior | National research institution | 27 | Constrained Comm |
| Sci_07 | France | ECR | National research institution | 2 | Constrained Comm |
| Sci_08 | US | Senior | Corporate research institution | 4 | Media Enthusiast |
| Sci_09 | US | ECR | University | 5 | Strategist |
| Sci_10 | US | Mid-career | University | 10 | Media Enthusiast |
| Sci_11 | US | Mid-career | University | 4 | Ambivalent Media S |
| Sci_12 | US | Senior | University | 5 | Media Enthusiast |
| Sci_13 | US | Mid-career | Nonprofit research institution | 10 | Media Enthusiast |
| Sci_14 | US | Mid-career | University | 15 | Ambivalent Media S |
| Sci_15 | Canada | Senior | University | 15 | Strategist |
| Sci_16 | US | Mid-career | University | 8 | Media Enthusiast |
| Sci_17 | England | Mid-career | Public health center | 5 | Strategist |
| Sci_18 | US | ECR | University | 10 | Strategist |
| Sci_19 | US | Mid-career | University | 1 | Media Enthusiast |

## Acknowledgment

The authors thank Kaylee Fagan for her work as a research assistant.

## Footnotes

1 Full data set and description of collection methods available on Github at [anonymized for review]

2 Both protocols are publicly available at [anonymized for peer review].

